# New-to-nature PHA synthase design using deep learning

**DOI:** 10.1101/2024.10.09.616406

**Authors:** Tuula Tenkanen, Anna Ylinen, Paula Jouhten, Merja Penttilä, Sandra Castillo

## Abstract

Polyhydroxyalkanaoate (PHA) synthases are a group of complex, dimeric enzymes which catalyse polymerization of Rhydroxyacids into PHAs. PHA properties depend on their monomer composition but enzymes found in nature have narrow specificities to certain R-hydroxyacids. In this study, a conditional variational autoencoder was used for the first time to design new-to-nature PHA synthases. The model was trained with natural protein sequences obtained from Uniprot and was used for the creation of approximately 10 000 new PHA synthase enzymes. Out of these, 16 sequences were selected for *in vivo* validation. The selection criteria included the presence of conserved residues such as catalytic amino acids and amino acids in the dimer interface and structural features like the number of *α*-helixes in the N-terminal part of the enzyme. Two of the new-to-nature PHA synthases that had substantial numbers of amino acid substitutions (87 and 98) with respect to the most similar native enzymes were confirmed active and produced poly(hydroxybutyrate) (PHB) when expressed in yeast *S. cerevisiae*. PHA including PHB have high potential as biodegradable and biocompatible materials. Ultimately the model-designed new-to-nature PHA synthases, could expand the PHA material properties to suit new application areas.

## Introduction

Polyhydroxyalkanoates (PHAs) are a group of polymers synthesized by several different bacteria (1). These bacteria synthesize PHAs as their carbon and energy storage under nutrient limited conditions when an excess of carbon source is present (2). PHAs have gained attention due to their excellent biodegradability and biocompatibility, making them environmentally friendly alternatives to conventional, nonbiodegradable thermoplastic materials.

One of the main challenges in replacing conventional plastics with PHAs is the brittleness of common commercial PHA grades, especially poly(hydroxybutyrate) (PHB) homopolymer and copolymers with high 3-hydroxybutyrate (3HB) content. This fragility is mainly caused by the growth of spherulites (higher-density spherical regions) during the storage of the final material, a phenomenon called secondary crystallization (3). Fortunately, the crystallinity degree can be reduced e.g., by an inclusion of other hydroxyacid monomers into the polymer chain. Similarly other material properties such as flexibility, permeability, and visual and thermal properties can be adjusted (1)(4)(5). As an example, poly(4hydroxybutyric acid) (4HB), a medical grade commercial PHA, is more stretchable, with a elongation at break value of 1000 % (6), and has lower glass transition temperature of -45 °C (7), in comparison to PHB, with corresponding values of 6 % and 3 °C.

The final monomer structure of the PHA chain depends on the availability of the corresponding coenzymeA (CoA) activated hydroxyacids and the substrate specificity of the PHA synthase enzyme which catalyzes the polymerization reaction. Thus, the PHA synthase has an important role in design of new polymer structures and properties. PHA synthases have been divided to four different classes based on their subunit structure and preference of monomers. Class I, III and IV PHA synthases prefer monomers with only three to five carbons (i.e., short-chain-length PHA monomers (SCL-monomers)), while class II PHA synthases prefer mediumchain-length PHA monomers (MCL-monomers) containing six to 14 carbons. Also the subunit structure differs between the classes. Class I and II PHA synthases are homodimers containing only PhaC subunits while class III and IV PHA synthases are heterodimers which have in addition to PhaC subunit a PhaE or a PhaR subunit, respectively. Active PHA synthases are dimers although recently Assefa et al. (2022) suggested that class I polyhydroxyalkanoate synthase from *Brevundimonas sp*. KH11J01 is active as a trimer (8).

By designing novel PHA synthases, it could be possible to produce PHAs with different monomer compositions and structures, thus tailoring their material properties for various applications. For example, protein design could be used to create PHA synthases that can incorporate both SCL- and MCL-monomers into the same polymer chain, resulting in PHAs with improved flexibility and toughness. Protein design could also be used to expand the range of monomers that PHA synthases can utilize, such as aromatic or halogenated monomers, which can introduce new functionalities and properties to PHAs. However, rational design of PHA synthases can be challenging. Crystal structures of the Cterminal catalytic domain has been resolved for only two PHA synthases, PHA synthases from *Cupriavidus necator* (9, 10) and *Chromobacterium* sp. USM2 (11). Furthermore, these crystal structures do not contain the N-terminal domain. In addition, several different catalytical mechanisms have been proposed (9, 10, 12–14), but no consensus on the mechanism exists. Instead of rational design, computational tools can be used. Computational protein design tools such as FuncLib (15) and CADENZ (16) can be used to generate libraries with enzyme variants. Enzymes with enhanced activity or even new substrate specificities has been found by screening these libraries (17). In addition, to these tools generative AI models, such as variational autoencoders (VAEs), can be used to generate libraries with new-to-nature enzymes. While VAEs have been successfully applied to design new-to-nature proteins such as metalloproteins, luciferase enzymes, and simpler proteins such as human-like phenylalanine hydroxylases (18) (19) (20), their potential for creating more complex enzymes such as dimers, like PHA syntases, has been understudied.

In this work, we designed two new-to-nature class I PHA synthase sequences and demonstrated their activity by polymerization of 3HB monomers in our previously developed yeast Saccharomyces *cerevisiae* strain expressing an 3HBCoA synthesizing pathway (21). New PHA synthase structures were designed using a generative AI model, specifically a conditional variational autoencoder (cVAE). New-tonature PHA synthases are interesting as they could expand the possibilities to polymerize different PHA monomers and adjust PHA material properties into new application areas. For example differences in substrate specificity could allow formation of block copolymer structures (structures formed by two or more different homopolymers) as shown earlier for chimeric PHA synthase PhaC_AR_ (22).

## Results

### A. Choice of a deep learning model

We trained an autoencoder model that takes as input the structural and sequence-based features of enzymes and outputs the amino acid sequences of enzymes. The model also receives a condition vector that represents the enzyme class. We used a dataset of PHA synthases, lipases, and partial PHA synthase sequences for the model training. We found that using bidirectional LSTM layers and multi-head self-attention blocks improved the model performance.

We defined the loss function of the model as a combination of the reconstruction loss and the KL divergence (23), which measures how much the latent space differs from the standard normal distribution. The KL divergence term can be scaled by a factor called beta that controls the trade-off between reconstruction and disentanglement of the latent features (24). However, a high beta value can lead to a posterior collapse, a phenomenon where the latent variables become independent of the input and the decoder ignores them. To mitigate this problem, we trained the model in stages, gradually increasing the beta parameter from 0.01 to 1.

We evaluated the performance of the model by three criteria: the coherence of the generated sequences with their corresponding PHA synthase class, amino acid composition and presence of ‘X’ inside the generated sequences (we used ‘X’ to mark the end of the sequence in the input sequences, which led the model to incorporate it also inside some generated sequences). The evaluation was performed by generating sequences in all four PHA synthase classes and analyzing them. The model produced sequences of correct class with high accuracy (93-100 %) but had more difficulty in matching the right amino acid composition (28-39 %) and avoiding ‘X’ inside the sequences (34-72 %) (Table 1). The model performed better in classes I and II than in classes III and IV in terms of amino acid composition and ‘X’ avoidance but had similar accuracy in class assignment across all classes.

**Table 1.** Evaluation of model performance. New-to-nature sequences were generated in all four PHA synthase classes (columns in the table). The first row “PHA synthase class” shows how many of the generated sequences were from correct class, the second row “Amino acid composition” shows percentage of sequences containing similar amount of each amino acid as the native PHA synthases and the third row “Presence of ‘X’” shows percentage of sequences containing ‘X’ inside the sequence.

|  | class I | class II | class III | class IV |
| --- | --- | --- | --- | --- |
| PHA synthase class | 100 % | 100 % | 93 % | 97 % |
| Amino acid composition | 33 % | 37 % | 39 % | 28 % |
| Presence of 'X' | 72 % | 63 % | 34 % | 66 % |

### B. Selection of 16 new-to-nature PHA synthase structures

We used our cVAE model to create over 10 000 PHA synthase sequences. As it is currently challenging, if not impossible, to experimentally characterize such a high number of different protein sequences, we carried out several filtration steps to select 16 most interesting sequences for an *in vivo* activity test. These steps included removal of duplicates, clustering based on similarity, and selection of only those that had the catalytic amino acids (i.e., Cys, Asp, His) in the active site and other conserved residues, including hydrophobic amino acids at the dimer interface. We also predicted the structures of the designed sequences with Alphafold (25), compared them with a PHA synthase from *Chromobacterium* sp. USM2 and checked the number of αhelixes in the N-terminal. In addition we analyzed the length of the sequences, tunnels to the active sites, and similarity with the closest natural enzymes. After these filtration steps, the resulting 16 selected enzymes (Table 2) showed highest sequence similarity to PHA synthases either from *Legionella* or *Janthinobacterium* species. To assess whether this bias was a result of the selection process or if all generated sequences were similar to these species, we constructed a phylogenetic tree from the initially generated sequences (Figure 2). The sequences generated using the model had remarkably higher variability than the sequences selected for *in vivo* tests showing its ability to avoid the usual issue of VAEs that reduce variance in the sampling (26).

**Table 2.**
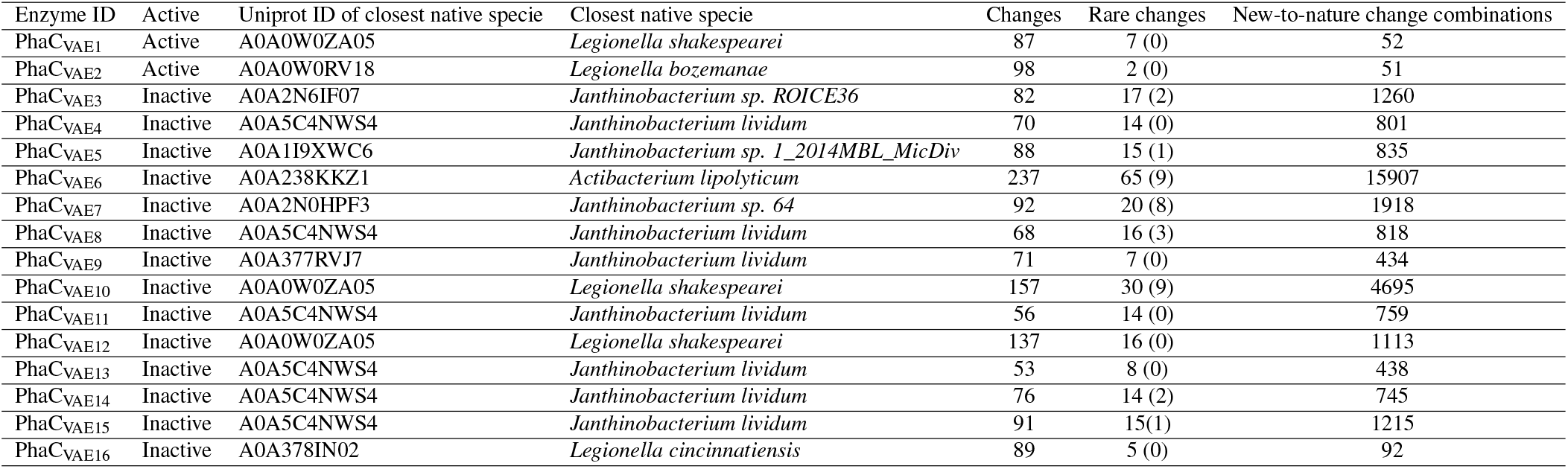
Generated enzymes. List of the generated PHA synthase enzymes using our cVAE model. The “active” column shows if the enzyme showed activity for 3HB-CoA polymerization when expressed *in vivo* in yeast *S. cerevisiae*. The second column is the Uniprot ID of the most similar natural enzyme and the third column is the organism where the most similar natural enzyme was found. The “Changes” column shows the number of different amino acids of the generated proteins compared with their most similar natural enzyme. The “Rare changes” column shows the number or amino acids present in less than 100 natural enzymes in the same position. Inside the parenthesis we show the number of new-to-nature changes. The column “New-to-nature change combinations” shows the number of two change combinations that were not present in any natural PHA synthase of class I.

**Table 3.** PHA synthase sequences used in training were divided to three different size categories based on their length. The table presents the amount of amino acids (aa) in the different categories for the four different PHA synthase classes. The first column presents the class of the PHA synthase.

| Class | Normal size | Small | Fraction |
| --- | --- | --- | --- |
| I | 500-700 aa | 200-499 aa | 0-199 aa |
| II | 450-650 aa | 200-499 aa | 0-199 aa |
| III | 300-500 aa | 200-299 aa | 0-199 aa |
| IV | 300-500 aa | 200-299 aa | 0-199 aa |

### *C. In vivo* activity test of the 16 new-to-nature PHA synthases

Together with PhaC_Ls_, PhaC_Jl_, and PhaC1_Cn_ encoding genes, the genes encoding for the selected 16 cVAE generated new-to-nature PHA synthases were individually integrated into chromosome X of *S. cerevisiae*, more specifically into X-4 EasyClone loci *(27)*. Parent strain contained three copies of the 3-hydroxybutyryl-CoA (3HB-CoA) pathway, including acetyl-CoA acetyltransferase (PhaA) and acetoacetyl-CoA reductase (PhaB1) (Table 6). The activities of the enzymes were first assessed by staining with Nile red that binds on the surface of the PHA granules (28). The Nile red staining suggested that two of the new-to-nature PHA synthases were active and polymerized 3HB monomers (Supplemental Figure 1A). The strains expressing PHA_VAE1 and PHA_VAE2 showed significantly higher based on twotailed paired Student’s t-Test (p<0.003) fluorescence (26.4 ± 1.6 Relative fluorescence units (RFU) and 59.8 ± 11.1 RFU, respectively) than the PHA negative control strain PhaA-PhaB1 (10.4 ± 1.0 RFU). The nonzero fluorescence intensity of the negative control is explained by the binding of Nile red to other intracellular structures such as lipids droplets (29). For comparison, positive controls PHA_Ls, PHA_Cn, and PHB_ctr2 showed similar fluorescence intensities (38.8 ± 5.4 RFU, 31.3 ± 5.4 RFU, and 48.8 ± 12.1 RFU, respectively) as the strains expressing the new-to-nature PHA_VAE1 and PHA_VAE2.

Next, the activities of the 16 new-to-nature PHA synthases were assessed by growing the yeast strains in shake flasks for 72 h and analyzing PHB content from the lyophilized biomass with precise gas chromatography mass spectrometry (GC-MS) method. The analysis of the PHB content in the biomass confirmed the earlier observations from the Nile red staining (Supplementary Figure 1B). Strains expressing the new-to-nature PHA synthases, PHA_VAE1 and PHA_VAE2, accumulated 6.2 ± 0.1 % and 4.5 ± 1.7 % of PHA as % of CDW, respectively, demonstrating activity of the newto-nature PHA synthases. For comparison, PHB content in the biomass of the CEN.PK113-7D was below the detection limit and the strain expressing only PhaA-PhaB1 and the strains expressing the other new-to-nature PHA synthases (PHA_VAE3-PHA_VAE16) showed only a trace amount of 0.05-0.15 % of PHB in CDW. The trace amount results likely from methanolysis of an excess of non-polymerized 3-HBCoA formed by PhaB1. Strains with native PHA synthases from, *C. necator* (PHA_ctrl1_), *L. shakespearei* (PHA_Ls_) and *J. lividum* (PHA_Jl_) accumulated 6.7 ± 0.1 %, 9.8 ± 1.0 %, and 13.3 % PHB of CDW, respectively. As PHA titers are linked to amount of formed biomass, the cell growth was followed at 0h, 24h, and 72h (Supplemental Figure 2). During the first 24 h, almost all strains reached their highest OD_600_ the only exception being strain PHB_ctr1 which showed minor OD_600_ increase after 24 h.

### D. Sequence analysis of the 16 new-to-nature PHA synthases

To identify the features that distinguished the active and inactive new-to-nature PHA synthases, we performed a comparative analysis of the sequences. Most of the 16 enzymes tested experimentally were similar to the native PHA synthases from two species: *Legionella sp* and *Janthinobacterium sp* (see table 2). Only two enzymes showed activity *in vivo* with the selected substrate, and they were similar to the natural enzymes from *Legionella shakespearei* and *Legionella bozemanae*. They had 87 and 98 amino acid substitutions, respectively, compared to their closest natural enzyme. The exact location of the substitutions is visualized in figure 3. The other enzymes, which were not active *in vivo*, had 53 to 237 substitutions compared to the most similar natural enzyme.

Next, we evaluated the frequency and novelty of the amino acid substitutions in each enzyme. The active enzymes exhibited less rare substitutions (amino acids with a frequency of less than 100 in natural enzymes at the same position) and no novel substitutions (amino acids absent in natural enzymes at the same position). The mean number of rare substitutions for the active enzymes was 4.5, while for the inactive enzymes it was 18.3. We also analyzed the pairwise combinations of substitutions in each enzyme. The active enzymes had 52 and 51 novel combinations, respectively, while the inactive enzymes had from 92 to 15907 novel combinations (see table 2).

## Discussion

In this study we created 16 new-to-nature PHA synthases with a conditional variational autoencoder. We found that two of these could effectively polymerize 3HB-CoA into PHB with accumulation levels similar to previously obtained by Ylinen *et al*. when expressing *C. necator* PHA synthase (30). Similar levels have previously been reported also for other engineered *S. cerevisiae* strains carrying the PHB pathway from *C. necator* (5.2-9% of CDW) (31) (32) (29), indicating that our new-to-nature PHA synthases were as efficient in polymerizing 3HB-CoA as the PHA synthase from *C. necator* when expressed in yeast. However, since it is difficult to normalize the effect of other parameters on PHA accumulation *in vivo*, precise comparison of different enzymes is difficult. For example, monomer availability has great impact on PHA accumulation, as shown in studies where replacement of PhaB1 with an NADH dependent alternative and the use of xylose or cellobiose as the carbon source instead of glucose boosted the carbon flux towards 3HB-CoA and result in higher PHB accumulation levels of 14-21 % of CDW in *S. cerevisiae* (33), (34) (30). Despite this, the *in vivo* approach is a rather easy method for assessing the capability of a PHA synthase to polymerize a certain monomer in relevant conditions.

The results of the comparative analysis of the 16 new-to-nature PHA synthases sequences suggest that the number and type of amino acid substitutions with respect to the most similar natural sequence may affect their activity *in vivo*. The two active new-to-nature PHA synthases contained significantly lower number of novel combinations of substitutions than the new-to-nature inactive proteins, suggesting that some combinations may be detrimental for the enzyme activity. We believe that taking this feature into consideration when selecting enzymes for wet lab experiments in the future is important.

Furthermore, improving the training dataset and protein representation of the input sequences might lead to higher success rate in producing active enzymes. Generative AI models typically require large amount of data for effective training. In addition to PHA synthase sequences we added lipase sequences to the training dataset as lipases share some structural similarities with PHA synthases (35). The effect of adding lipases to the training dataset on the quality of the generative model is unclear. It is possible that some of the lipases have divergent structures that could interfere with the model’s ability to learn the features of PHA synthases. However, the number of PHA synthase sequences in public databases is increasing rapidly. Therefore, in the future the model could be trained exclusively with PHA synthase sequences.

In this work, we utilized contact maps predicted by trRosetta (36) to represent the 3D structures of the input sequences for the deep learning model. Contact maps predicted by trRosetta contain information about the distances and orientation of amino acid residues that are in contact with each other in the enzyme structure. Thus, these contact maps describe enzymes better than their protein sequences. Predicting contact maps with trRosetta is fast and can be used easily on a data set of thousands of enzymes. However, the accuracy and quality of the protein representation, and thus also model’s performance, could be enhanced in future by the new advances in protein structure prediction such as Alphafold (25) or RoseTTAFold (37).

In addition, future work on PHA synthase design could explore other types of generative models instead of VAE. The VAE are efficient generative models which have been used successfully in the past to create new-to-nature proteins (20)(19)(18). However, they often encounter various challenges during the training process, such as posterior collapse, vanishing gradients, over-fitting, and dataset bias, which tend to make the training of VAEs very difficult and unstable (38). We addressed the problem of posterior collapse by adopting a phase training approach increasing gradually the beta parameter in the loss function. However, problems like posterior collapse, loss of variability or bias could also be avoided by using alternative types of generative models, such as diffusion models (39).

A direct comparison of our approach with other VAEs used for protein design, such as those mentioned in the introduction, may not be appropriate. The success rate of AI models depends on several factors, such as the model architecture and training, the quality and quantity of the training data, the data representation, and the complexity of the target protein. PHA synthases are poorly understood proteins, for which we lack a complete crystal structure, the polymerization mechanism is unclear, and both subunits of the dimer are required for enzyme activity, although their roles are unknown. All these factors affect the performance and success rate of the method. Therefore, we did not attempt a direct comparison of our method with others, but we showed that our method is able to produce efficiently active new-to-nature enzymes despite the challenges and uncertainties in the protein design of PHA synthases. Our future goal is to use this model to design PHA synthases that can utilize a wider or more specific range of substrates, or that can produce novel polymers with desirable properties.

## Conclusions

In this study, a conditional variational autoencoder was used for the first time to create new-to-nature PHA synthases that are active as dimers. Despite 87 and 98 amino acid substitutions in comparison to the closest native PHA synthases the two PHA synthases were active and produced PHB in yeast *S. cerevisiae*. Ultimately these, or other new-to-nature structures designed in future, could expand possibilities to polymerize different PHA monomers and adjust PHA material properties into new application areas.

The success rate of our low-throughput approach was 12.5% for designing new-to-nature PHA synthases. This enzyme is very challenging to engineer, as it is poorly characterized, lacks a complete crystal structure, and requires dimerization for its function. The conditional variational autoenconder generated a diverse set of PHA synthases sequences by learning from natural proteins, and we applied a series of tests to select the most promising candidates. We believe that the selection process may have contributed to the high success rate. However, our subsequent analysis revealed that the low frequency of unnatural amino acid pair combinations was a key feature of the active sequences, and we suggest using this criterion for future selections.

## Materials and Methods

### E. Data collection

#### E.1 Collection and processing of PHA synthase data

We used sequences from Uniprot (40) (uniprot version 2021_03) to create a data set for this study. These included all sequences containing class I, II, or III PHA synthase domains (InterPro families IPR010963, IPR011287, IPR010125 (41)), or N-terminal domain of a PHA synthase (InterPro family IPR010941 (41)). A class was assigned for each sequence based on InterPro family and a phylogenetic tree. ClustalW 2.1 (42) was used to create a multiple sequence alignment (MSA) of sequences belonging to InterPro (41) classes IPR010963, IPR011287 and IPR010125. The D3 javascript library (43) was used to create a phylogenetic tree of the MSA. A cluster with known class IV poly(R)-hydroxyalkanoate synthase sequences (Q8GI81 and Q9ZF92, Uniprot) was assigned as class IV. The sequences assigned to class IV belonged to class III polyhydroxyalkanoate synthase family in InterPro (IPR010125). Thus, the sequences belonging to IPR010963 were assigned to class I, the sequences belonging to IPR011287 were assigned to class II, and the sequences belonging to IPR010125 were assigned to class III or IV based on the MSA and phylogenetic tree explained above. For sequences belonging to IPR010941 but not to IPR010963, IPR011287, or IPR010125, the class was assigned based on Basic Local Alignment Search Tool (BLAST). BLAST 2.12.0 (44) was run for each sequence against PhaC sequences belonging to IPR010963, IPR011287 or IPR010125. The class of the query sequence was assigned to be the same class as the PHA synthase sequence with lowest e-value. Furthermore, we augmented the data set with sequences with known beneficial mutations (Supplementary Table 1). In addition to the PHA synthase sequences, 5000 lipase sequences were included in the dataset. Lipases were added to increase the size of the dataset as they have similar structures as the PHA synthases. Both lipases and PHA synthases contain α/β-hydrolase domains. Furthermore, PHA synthases contain lipase box sequences (GXSXG) with only one amino acid substitution compared to lipases. In PHA synthases, the active site serine is replaced with cysteine (35). First, sequences containing lipase in their name were downloaded from Uniprot (v. 2021_3). Then BLAST 2.12.0 was run for these sequences against PhaC sequences belonging to IPR010963, IPR011287, or IPR010125, and 5000 lipase sequences with smallest e-value were included in the dataset. Next, the sequences were classified based on their size. Very long sequences were removed and the rest of the sequences were divided to three different size categories (i.e., normal size, small, and fraction) based on the sequence length (table 3). Finally, the dataset was divided to training and testing datasets. The CD-hit 4.6 (45) was used to cluster all sequences in each class (i.e., Class I-IV PHA synthases and lipases) separately. One cluster in each class was then selected to testing dataset. The data processing pipeline is clarified in Supplementary Figure 3. The number of sequences in different classes, the number of sequences in different size categories as well as the sizes of training and testing dataset are compiled in Table 4.

**Table 4.** Amount of sequences in different size categories, training dataset and testing dataset.

| Class | Total | Normal size | Small | Fraction | Training dataset | Testing dataset |
| --- | --- | --- | --- | --- | --- | --- |
| I | 10405 | 9696 | 432 | 277 | 10200 | 205 |
| II | 3128 | 2995 | 96 | 37 | 2867 | 261 |
| III | 1602 | 1536 | 12 | 54 | 1527 | 75 |
| IV | 500 | 469 | 14 | 17 | 472 | 28 |
| lipases | 5000 |  |  |  | 4516 | 484 |
| in total | 20635 | 14696 | 554 | 385 | 19582 | 1053 |

#### E.2 Protein representation

We used a combination of features to represent the sequence and structure information of the enzymes for the generative model. To obtain structural features for our large dataset of enzymes, we used the software trRosetta (46). It predicts the contact maps of each enzyme, which consist of a matrix of distance probabilities and three matrices of orientation probabilities for each pair of residues. Moreover, we encoded each protein sequence using seven physico-chemical amino acid properties, such as mass, side chain volume, or polarity (see table 5), and the predicted secondary structure of the protein. We evaluated the performance of different protein representations for various models, andfound that the combination of these three features (contact maps, amino acid physico-chemical features and secondary structure prediction) yielded the highest accuracy.

**Table 5.** Features collected from each amino acid.

| Property | Definition | Reference |
| --- | --- | --- |
| Mass | Masses of neutral residues | (47) |
| Side chain volume | Mean volumes of residues inside proteins | (48) |
| Hydropathy | Hydrophobicity/hydrophilicity of the residues | (49) |
| Polarity | Average separation of charge in the residues | (50) |
| Polarizability | Possibility of the residues to become polarized temporality | (51) |
| pI | pH of the residues at the isoelectric point. | (52) |
| Side chain composition | Atomic weight ratio of noncarbon elements inside side chain end groups or rings | (50) |

**Table 6.**
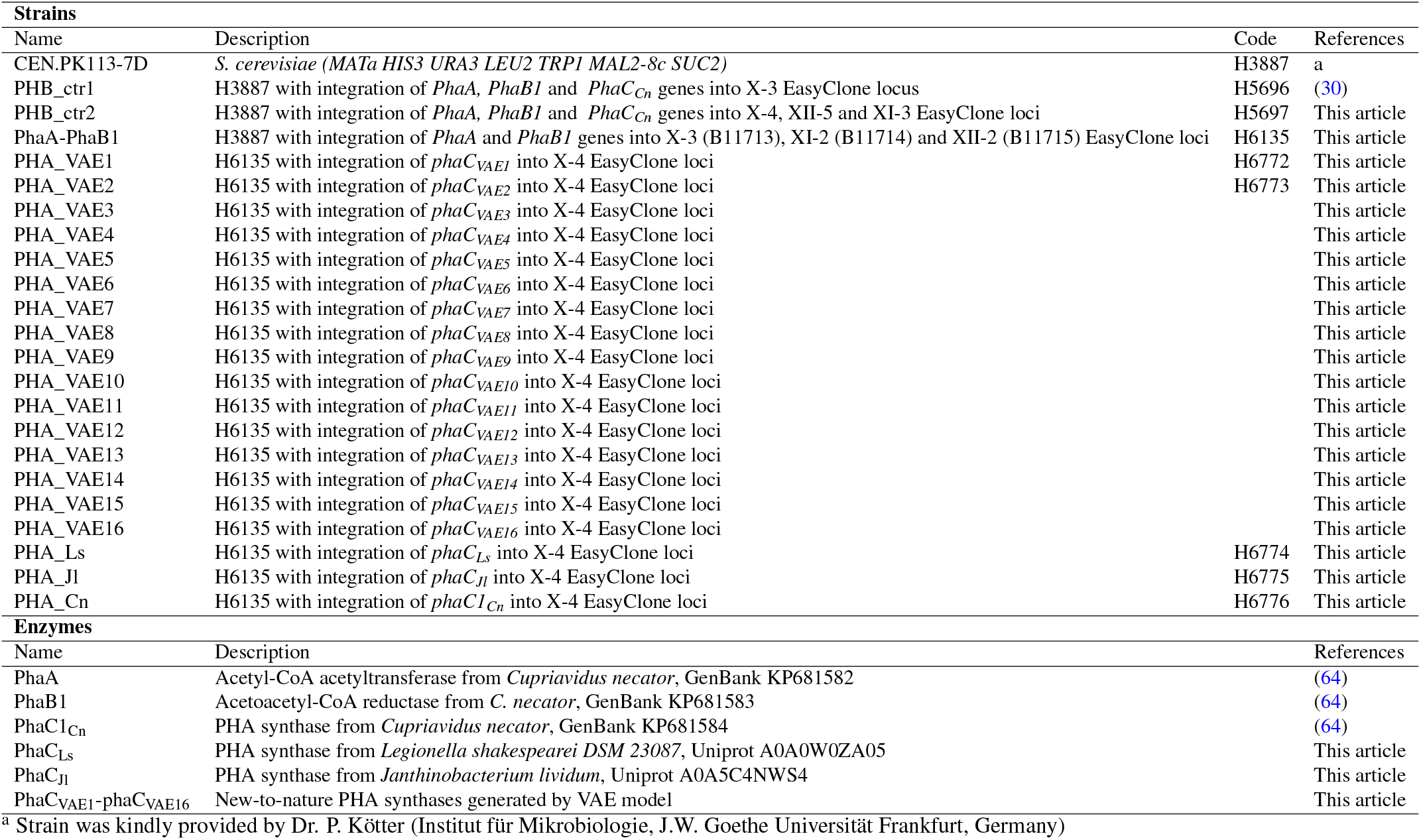
Strains and enzymes used in this study.

### F. Building and use of a variational autoencoder (VAE)

#### F.1 Model architecture

Our model follows an autoencoder architecture, which consists of two parts: the encoder that compresses the input into a latent vector and the decoder that reconstructs the input from the compressed latent vector created by the encoder. However, our model differs from the standard autoencoders because it does not use the same input and output. Our encoder takes as input the structural and sequence-based features of enzymes, while the decoder outputs the one-hot representation of the amino acids in the enzyme sequence. An additional input to both the encoder and the decoder is the condition that represents each enzyme class (a 85-dimensional vector). The enzyme data for the model training comprised PHA synthases (4 classes), lipases (1 class), and partial PHA synthase sequences (4 classes) (Table 4. We separated the partial sequences based on length and treated them as distinct classes in the condition vector.

We compared the convolutional (53) and recurrent (LSTM (54) and GRU (55)) model architectures and selected the LSTM-based model because it achieved the highest test accuracy. Autoencoder architectures consist of two parts: the encoder, which compresses the input given for the training, and the decoder, which reconstructs the input from compressed values produced by the encoder. In our model, the encoder input comprised the contact map representation of the protein (a 700×75 matrix), the protein amino acid features representation (a 700×7 matrix) (table 5, the secondary structure prediction (a 700×3 matrix), and the condition vector (an 85dimensional vector). Each of these inputs, except the condition, was passed through a multi-head attention block (56) with 18 heads, followed by a bidirectional LSTM layer. The outputs of these three blocks were then concatenated with the condition vector and fed into two dense layers to produce the mean and the logvar values corresponding to the input.

We sampled the values of the latent vector using the mean and the logvar values generated by the encoder. The decoder had two inputs: the latent vector sampled from the encoder (a 20-dimensional vector) and the same condition vector used in the encoder. Both inputs were concatenated and connected to a bidirectional LSTM layer. The output of the LSTM layer was flattened and sent to two consecutive dense layers that returned a one-hot representation of the amino acids in the protein sequence (a 700×21 matrix) (see figure 1)

**Fig. 1.**
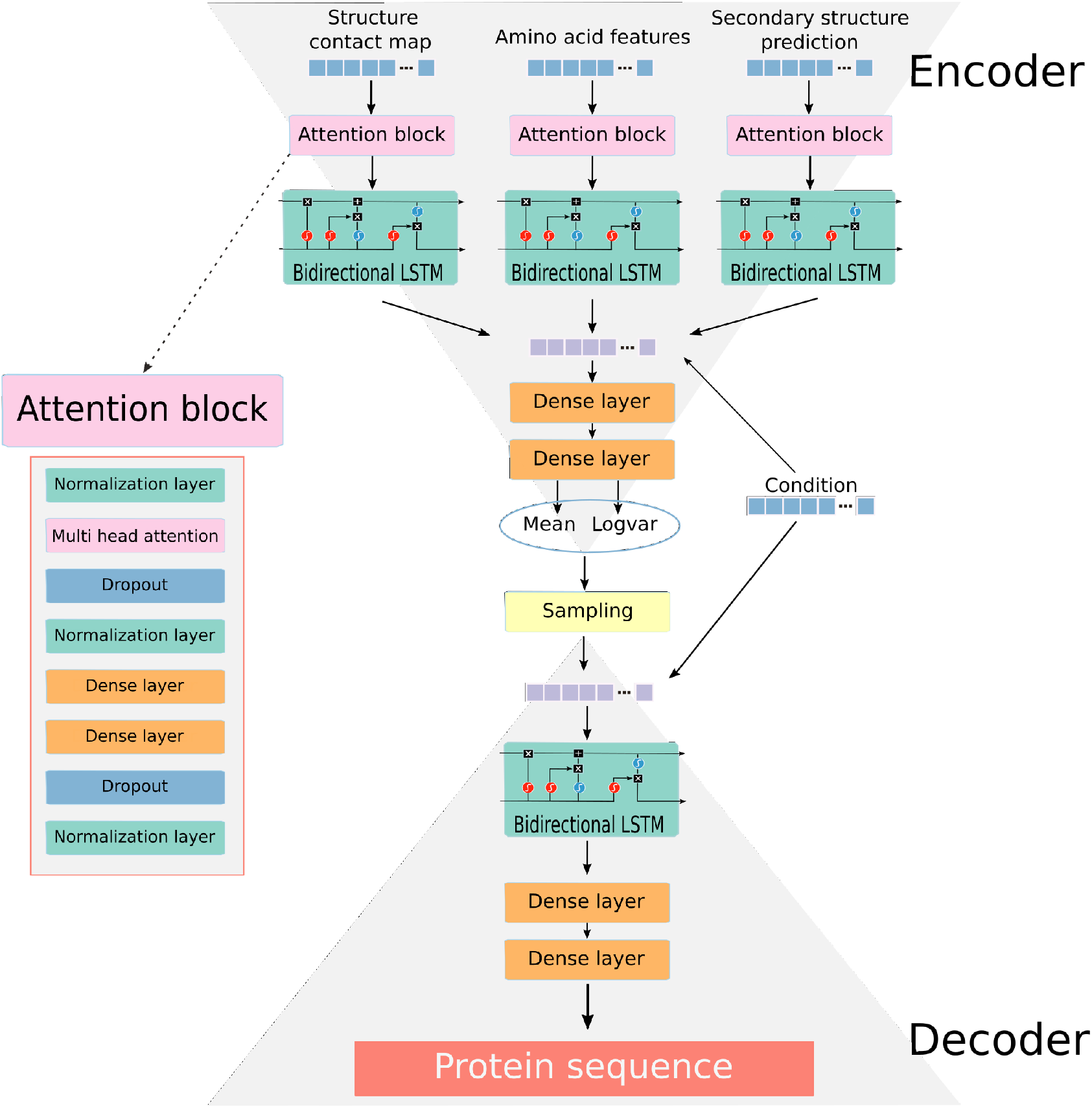
Scheme of the deep learning model used to generate new to nature enzymes.

**Fig. 2.**
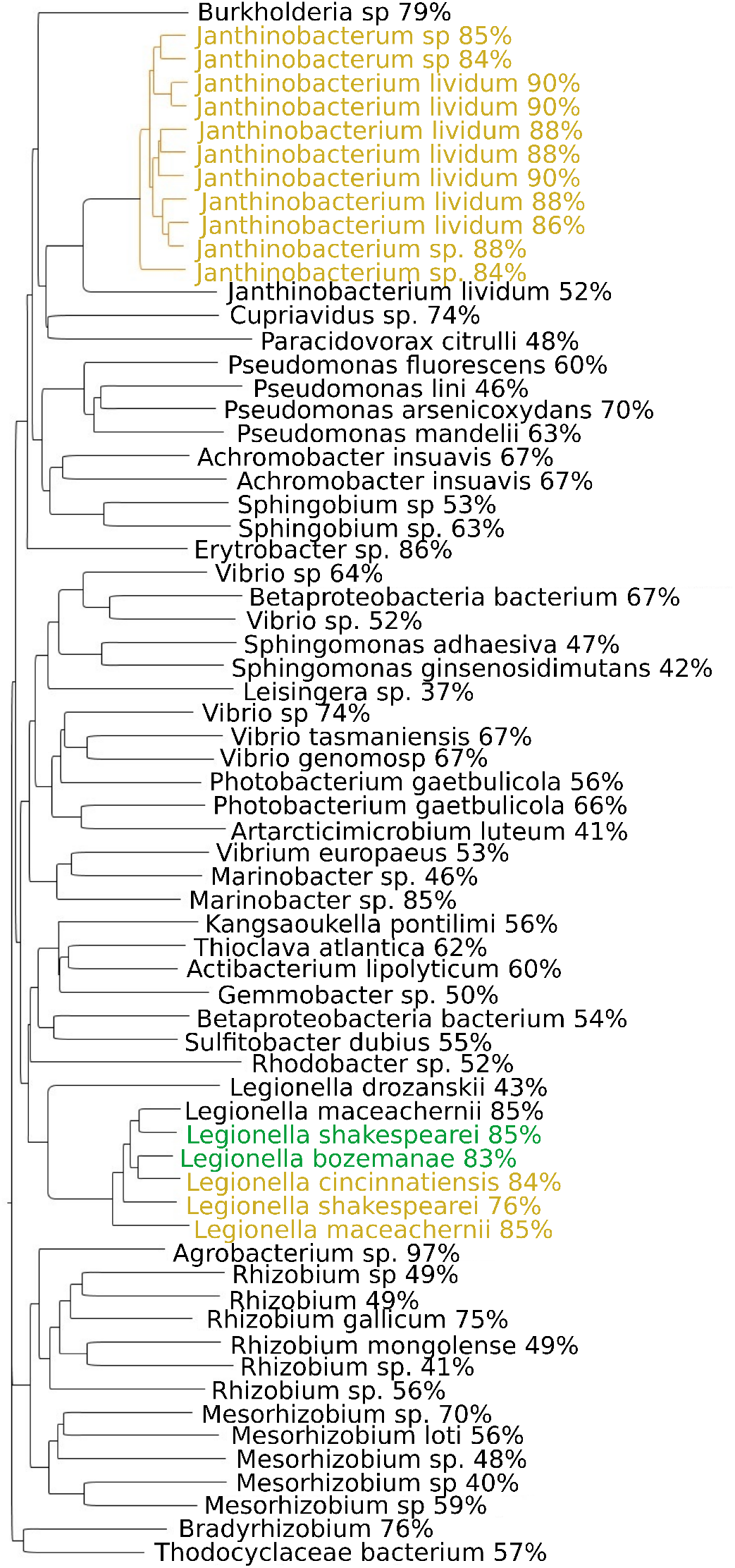
Phylogenetic tree of cVAE generated sequences. Labels in the tree describes the closest native PHA synthase and percentage identity between the sequences. Sequences selected for *in vivo* activity analysis are colored with green (active) and orange (not active).

**Fig. 3.**
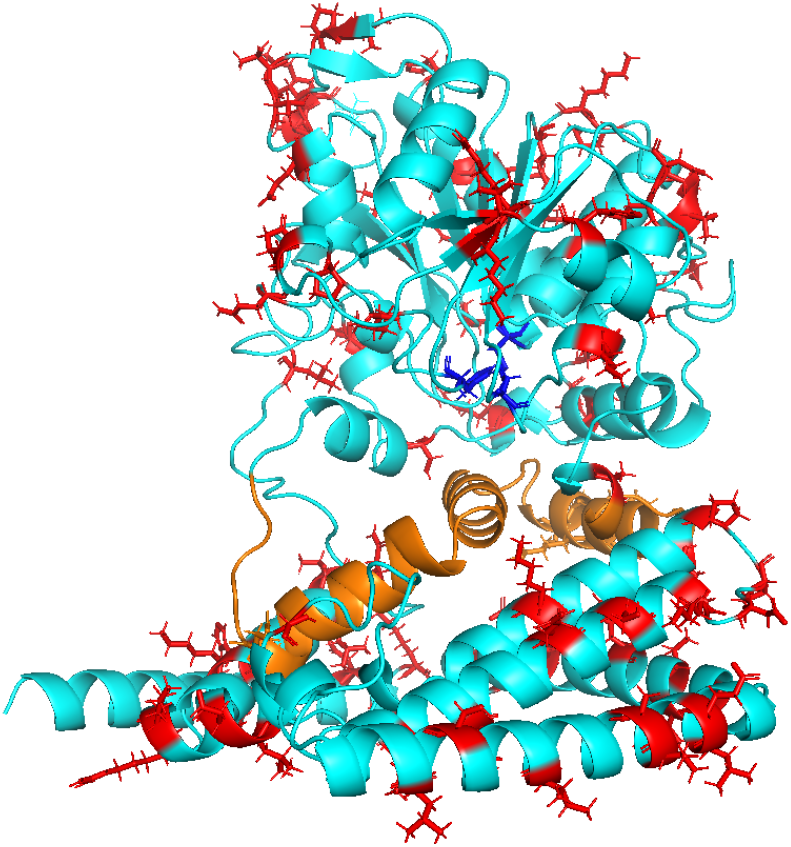
Structure of the generated active enzyme PhaCVAE2 predicted with Alphafold (25). The 98 different amino acids with respect the most similar natural enzyme are marked with red. Catalytic amino acids are shown with dark blue. The orange alpha helices are part of the CAP region of the enzyme.

#### F.2 Model training and evaluation

The model was trained using Adam optimizer with learning rate 0.001 and using batch size of 12. First training was done for approximately 200 epochs using beta parameter 0.01, then approximately 200 epochs using beta parameter 0.1 and finally for approximately 200 epochs using beta parameter 1. The beta parameter controls the weight of KL divergence on the total loss. Softmax cross entropy was used to calculate the reconstruction error in the loss function. Model performance was evaluated using categorical accuracy as well as analyzing quality of the produced sequences. First we evaluated if the model produced sequences in the correct class. This was done by generating 100 sequences in each class, running BLAST 2.12.0 against native PhaC sequences (belonging to InterPro families IPR010963, IPR011287 or IPR010125) and evaluating if the hit with the lowest e-value was from correct class. Next we analysed if the sequences produced had similar amount of each amino acid as the natural PHA synthases. We selected 1000 native sequences from each PHA synthase class randomly (InterPro families IPR010963, IPR011287, IPR010125) and counted the amount of each amino acid in these sequences. This way we defined the range of each amino acid in the native sequences (e.g., amount of alanine residues was 4-16% in the native sequences). We then generated 1000 new PHA synthase sequences from each class and analyzed how many of them had all the amino acids within the calculated range. Finally we analyzed if the model generated sequences containing “X” inside the sequence. As all inputs had to have length of 700 we added sequence of “X” in the end of each sequence shorter than 700 amino acids to obtain uniform length of 700 amino acids. This made the model to add “X” also in the middle of some generated sequences. However, as the aim was to have “X” only in the end of the sequence to mark the end of the enzyme the amount of sequences containing “X” also elsewhere was analyzed.

#### F.3 Sequence generation and selection of the 16 sequences

We generated 10322 different class I PHA synthase sequences using the selected cVAE generative model and by randomizing the values in the latent space. When generating sequences, only the sequences having amino acid proportions within the same range as in the native sequences (calculated similarly as in Materials and methods, Model training and evaluation) were saved. Next we removed duplicated sequences and filtered the generated PHA synthases based on the presence/absence of the three catalytic amino acids in the right position in active site leaving 9241 enzymes. With the remaining sequences, we used cd-hit (45) with 0.95 of similarity threshold to cluster them resulting on 715 clusters. When generating sequences cVAE gave the probability of each amino acid for each position in the sequence. The amino acid with highest probability was then selected to the generated sequence. To select sequences from the clusters we analysed for each position in the sequence, how many amino acids had similar probability as the selected amino acid in each position of the sequence(maximum of 10 % smaller probability than the amino acid selected to the sequence). Then from each cluster we selected the sequence with least of amino acids having similar probabilities than the amino acid with highest probability as we considered that the model was less confident in those positions. Next the filtering continued by running AlphaFold 2 (25) for each of the 715 sequences and aligning the obtained structures with the crystal structure of PHA synthase from *Chromobacterium* sp. USM2 (PDBe: 5xav (35)) using TM-align (57). These aligned structures were then used to analyze if the created new PHA synthase sequences had changes in conserved residues (other than the catalytic triad) obtained from Check *et al*. (35) (amino acids corresponding to 197L, 200Y, 211P, 213L, 220N, 223Y, 226D, 232S, 249W, 289G, 291C, 293G, 294G, 323D, 365R, 392W, 395D, 415N, 431D, 448H, 476G, 489K in *Chromobacterium* USM2). In addition, we studied if amino acids in the dimer interface corresponding to amino acids at positions 332, 333, 369, 371, 386, 387, 390, and 451 in *Chromobacterium* sp. USM2 (35) were hydrophobic. However, the alignment of our structures with 5xav was not good for positions 371 and 386 in PHA synthase of *Chromobacterium* sp. USM2. We expect this to be due to the break in the crystal structure at positions 372-384. Thus, we aligned the generated 715 sequences also with the AlphaFold structure of the PHA synthase from *Chromobacterium* sp. USM2 (AlphaFoldDB: E1APK1) and used these alignments to check the hydrophobicity of the amino acids corresponding to amino acids at positions 372-384 of E1APK1. Next we analyzed the amount of α-helixes in the N-terminal part of the cVAE generated PHA synthase sequences. Kim *et al*. (58) analyzed that five α-helixes in the N-terminal are required for the enzyme to function properly. Thus, we analyzed the amount of α-helixes in the Nterminal of our new sequences by aligning AlphaFold predicted structures with PHA synthase from textitCupriavidus necator (AlphaFoldDB: P23608), analyzing the start of the N-terminal from the aligned structures and predicting the secondary structure from the AlphaFold predicted structures using STRIDE (59). Then, we used Miyata’s distance (60) to compare the amino acid replacements between each generated sequence and the PHA synthase from *Chromobacterium* sp. USM2 and Cupriavidus necator. We divide the enzymes in different areas: the catalytic sites, the amino acids around the catalytic sites in the active pocket, the CAP, the N-terminal part and the rest of the enzyme. Miyata’s distance was calculated as the sum of the square distance between the amino acids polarity and volume divided by the standard deviations. The score was then normalized by the number of changes and the length of the specific area. The distance of the enzymes on critical areas was used as one of the criteria for the final selection of the enzymes to be tested experimentally. Furthermore, we checked the lengths of the generated sequences and amount of ‘X’ inside the sequence similarly as when analyzing the models (Materials and methods, Sequence generation and selection). For sequences containing ‘X’ inside the sequence, the ‘X’ was changed to the amino acid with second highest probability. With the information explained above, we then selected 42 sequences for further analysis. For these 42 sequences, we calculated the tunnels to and from the active site with CAVER 3.0.3 Pymol plugin (61) and finally selected 16 to be tested in wet lab. To analyze if the selection process led us to select sequences uniformly from all the produced sequences, we generated a phylogenetic tree of the cVAE generated sequences. First, we clustered all class I PHA synthase sequences generated with the model using CD-hit (45). This resulted in one representative sequence for each cluster. Next, all 16 sequences selected for *in vivo* experiments were added to the set of sequences. BLAST 2.15.0 (44) was then run against native PHA synthase sequences (belonging to InterPro families IPR010963, IPR011287 or IPR010125) and the hit with lowest percentage identity was selected. A phylogenetic tree of the cVAE generated sequences was then generated using ClustalW 2.1 (42) and visualised using Geneious 10.2.6 (https://www.geneious.com). The closest native sequences were marked in the branch labels.

### *G. In vivo* activity measurement of new-to-nature PHA syntases

#### G.1 Strain engineering

Strains and enzymes used in this study are presented in Table 6. The used plasmids are listed in Supplementary Table 2. *S. cerevisiae* CEN.PK113-7D was kindly provided by Dr. P. Kötter (Institut für Mikrobiologie, J.W. Goethe Universität Frankfurt, Germany). Pathway to produce 3-HB-CoA (*phaA* and *phaB1*) was cloned to three different EasyClone vectors pCfB3034, pCfB2903, and pCfB3039 by amplifying the precursor pathway (pTEF1-phaA-tENO1-pTDH3-phaB) from plasmid pPHB_template_1_ (B9660) (62) and cloning the product into the corresponding EasyClone plasmids in front of *tCYC1* terminator using Gibson assembly (E2611S, New England BioLabs). The generated plasmids were digested with NotI and transformed to CEN.PK113-7D (H3887) to generate strain PhaA-PhaB1 (H6135). Gene coding for phaC1_Cn_ (*pPGK1phaC1*_*Cn*_) was amplified with PCR from pPHB_template_1_ (B9660) and cloned to EasyClone vector pCfB3035 in front of *tCYC1* terminator with Gibson assembly. Genes coding for VAE generated new-to-nature PHA synthases (PHAC_VAE1_PHAC_VAE16_) and genes coding for PhaC_Ls_ and PhaC_Jl_ were codon optimized for *S. cerevisiae* and ordered with PGK1 promoters from GenScript in pCfB3035 EasyClone vectors. The plasmids were digested with NotI and transformed to parent strain PhaA-PhaB1 (H6135) to generate strains PHA_VAE1-VAE16. Strain PHB_ctr2 was built by amplifying PHB pathway (pTEF1-phaA-tENO1-pTDH3phaB1-tSSA1-pPGK1-phaC1_Cn_-tCYC) with PCR from the pPHB_template_2_ (B11787) and cloning the product to EasyClone vectors pCfB3035, pCfB2904, and pCfB2909 using Gibson assembly. These vectors were then digested with NotI and transformed to CEN.PK113-7D (H3887). Lithium acetate (LiAc)/ SS carrier DNA/ PEG method (63) and CRISPR/Cas9 protocol of the EasyClone kit (27) were used in all transformations. Correct integrations were confirmed with PCR using oligos of EasyClone kit and gene specific oligos as well as Sanger sequencing (Microsynth Seqlab GmbH).

#### G.2 Nile red analysis

The strains were grown for 16 h in 3 ml of synthetic complete media with 20 g/l of glucose in a 24 well plate at 770 rpm shaking and 30 °C. Cultivation was started by inoculating the media with the strains from YPD plates. At the end of the cultivation each cell culture was diluted with distilled water to obtain OD_600_ 2. Then 100 µl of each of the diluted samples was transferred to a black 96-well plate and mixed with 20 µl of Nile red dissolved in DMSO, so that the final concentrations of Nile red and DMSO were 5 mg/l and 17 % v/v, respectively. Fluorescence was measured after 10 min of incubation at RT with Varioskan Flash (Thermo scientific) using 550 nm excitation and 610 nm emission wavelengths. Each culture was measured in four technical replicates.

#### G.3 Shake flask cultivation

All the strains were grown in synthetic complete media with 20 g/l of glucose at 30 °C and 220 rpm shaking. Precultures of 10 ml in 50 ml Erlenmeyer flasks were grown overnight. Subsequently 50 ml cultures in 250 ml Erlenmeyer flasks were started from OD_600_ 0.2 and continued for 72 hours. Samples were taken at 24 h and 72 h to monitor the population growth, extracellular metabolite formation, and glucose utilization. At the end of the 72 h cultivation, cells were harvested by centrifuging them for 6 min at 4000 rpm and washing once with distilled water. However, the washing step was not done for strains PHA_VAE3_A-C_, PHA_VAE4_A-C_, PHA_VAE5_A-B_, PHA_VAE6_A_, PHA_VAE7_A_ and PHA_VAE9_A-B_. Here A, B, and C refers to three different biological replicates of each strain. The cell pellets were stored in -20°C until GC-MS analysis. The population growth was analyzed by measuring the optical density (OD_600_).

#### G.4 PHB quantitation with GC-MS

The amount of accumulated PHB in CDW was analyzed with gas chromatography mass spectrometry (GC-MS) similarly to Ylinen *et al*. (30) based on method described by Braunegg *et al*. (65). Cell samples from the shake flask cultivations were frozen at -80 °C and lyophilized overnight in Christ Alpha 2-4 LSCBasic device. Ten milligrams of each dried sample was subjected to methanolysis for 140 min in 100 °C water bath in a solution containing 1 ml chloroform, 810 µl methanol, 150 µl sulfuric acid, and 50 µl 3-hydroxybutyric acid 1,3-13C2 (SigmaAldrich) as an internal standard. Samples were cooled down to room temperature and 1 ml of distilled water was added to remove water-soluble particles. Chloroform phase was then analyzed using gas chromatography equipment (7890, Agilent) with HP-FFAP column (19091F-102 Agilent). Two replicates were analyzed of each strain. A 3-hydroxybutyric acid standard was analyzed equally as the samples.

### H. Sequence analysis of the generated new-to-nature enzymes

Generated protein sequences were blasted against all the PHA synthase sequences collected from Uniprot (40) to identify the most similar natural enzyme for each novel protein. Then generated enzyme were structurally aligned to their most similar natural PHA synthase enzymes using TMalign (66) and their amino acid differences were recorded. To collect the new-to-nature change combinations, an additional structural alignment using TMalign with the *Cupriavidus necator* class I PHA synthase was performed. Then each position where a different amino acid was found in the generated enzyme was compared with the most similar natural enzyme. All the natural enzymes with the same amino acid in the same position were collected. If the number of collected natural enzymes was less than one hundred we considered the difference at that position as a rare change. From the collected natural enzymes, we then checked the location where the rest of the changes were and counted the cases where there was no natural enzyme with the combination of two changes in their corresponding places.(see table 2)

## Supplementary Note 1: Acknowledgement

We would like to express our gratitude to the Jane and Aatos Erkko Foundation (JAES) under project 220048 (Virtual laboratory for Biodesign, JAES-BIODESIGN) and the Wihuri Foundation for their generous support in covering the expenses associated with this work. Their contributions have been invaluable in enabling us to conduct this research. Additionally, we would like to acknowledge the invaluable support of Kaisa Peltonen in the lab and Samuli Ollila for comments on the manuscript. Their assistance has been crucial to our efforts.

