## Supplemental table 1 for "New-to-nature PHA synthase design using deep learning"

Supplementary Table 1. PHA synthases with known good mutations added to the training dataset.

| Uniprot ID | organism | mutations | class | Reference |
| --- | --- | --- | --- | --- |
| EIAPK1 | Chromobacterium sp. USM2 | A479V | I | Chuah et al., 2013 |
| EIAPK1 | Chromobacterium sp. USM2 | A479T | I | Chuah et al., 2013 |
| EIAPK1 | Chromobacterium sp. USM2 | A479Q | I | Chuah et al., 2013 |
| EIAPK1 | Chromobacterium sp. USM2 | A479M | I | Chuah et al., 2013 |
| EIAPK1 | Chromobacterium sp. USM2 | A479G | I | Chuah et al., 2013 |
| G0ETI5 | Cupriavidus necator | A510M | I | Tsuge et al, 2004 |
| G0ETI5 | Cupriavidus necator | A510E | I | Tsuge et al, 2004 |
| G0ETI5 | Cupriavidus necator | A510D | I | Tsuge et al, 2004 |
| G0ETI5 | Cupriavidus necator | A510C | I | Tsuge et al, 2004 |
| G0ETI5, O32471 | Cupriavidus necator, Aeromonas caviae | * | I | Matsumoto et al., 2009 |
| G0ETI5, O32471 | Cupriavidus necator, Aeromonas caviae | *, N149D, F314L | I | Phan et al., 2022 |
| O32471 | Aeromonas caviae | N149S, D171G | I | Tsuge et al., 2007 |
| O32471 | Aeromonas caviae | N149S, D171G, S389T | I | Harada et al, 2021 |
| O32471 | Aeromonas caviae | F518I | I | Amara et al, 2002 |
| O32471 | Aeromonas caviae | F362I, F518I | I | Amara et al, 2002 |
| O32471 | Aeromonas caviae | D459V, A513C | I | Amara et al, 2002 |
| Q848R9 | Pseudomonas stutzeri | S326T, Q482K | II | Shen et al, 2011 |
| Q9Z3Y1 | Pseudomonas sp. 61-3 | S325T | II | Takase et al, 2003 |
| Q9Z3Y1 | Pseudomonas sp. 61-3 | S325C | II | Takase et al, 2003 |
| Q9Z3Y1 | Pseudomonas sp. 61-3 | Q481R | II | Takase et al, 2003 |
| Q9Z3Y1 | Pseudomonas sp. 61-3 | Q481M | II | Takase et al, 2003 |
| Q9Z3Y1 | Pseudomonas sp. 61-3 | Q481K | II | Takase et al, 2003 |
| Q9Z3Y1 | Pseudomonas sp. 61-3 | E130D, S325T, S477G, Q481K | II | Yang et al., 2011 |
| Q9X5X7 | Pseudomonas resinovorans | E130D, S325T, S477G, Q481K | II | Yang et al., 2011 |
| G3XCV5 | Pseudomonas aeruginosa | E130D, S325T, S477G, Q481K | II | Yang et al., 2011 |
| C0LD26 | Pseudomonas chlororaphis | E130D, S325T, S477G, Q481K | II | Yang et al., 2011 |
| B9W0T0 | Pseudomonas sp. MBEL 6-19 | E130D, S477H, Q481M | II | Yang et al., 2011 |
| B9W0T0 | Pseudomonas sp. MBEL 6-19 | E310D, S325T, S477R, Q481M | II | Yang et al., 2011 |
| B9W0T0 | Pseudomonas sp. MBEL 6-19 | E310D, S325T, S477G, Q481K | II | Yang et al., 2011 |
| B9W0T0 | Pseudomonas sp. MBEL 6-19 | E310D, S325T, S477F, Q481K | II | Yang et al., 2011 |
| B9W0T0 | Pseudomonas sp. MBEL 6-19 | E310D, S325T, Q481K | II | Yang et al., 2011 |
| B9W0T0 | Pseudomonas sp. MBEL 6-19 | E310D, Q481K | II | Yang et al., 2011 |

*25 % of N-terminal region of PhaC_Ac_ (O32471) and 74 % of N-terminal region of PhaC_CN_ (G0ETI5).
