## Supplementary figures and images for "New-to-nature PHA synthase design using deep learning"

### Supplemental table 2

Supplementary Table 2. Plasmids used in this study.

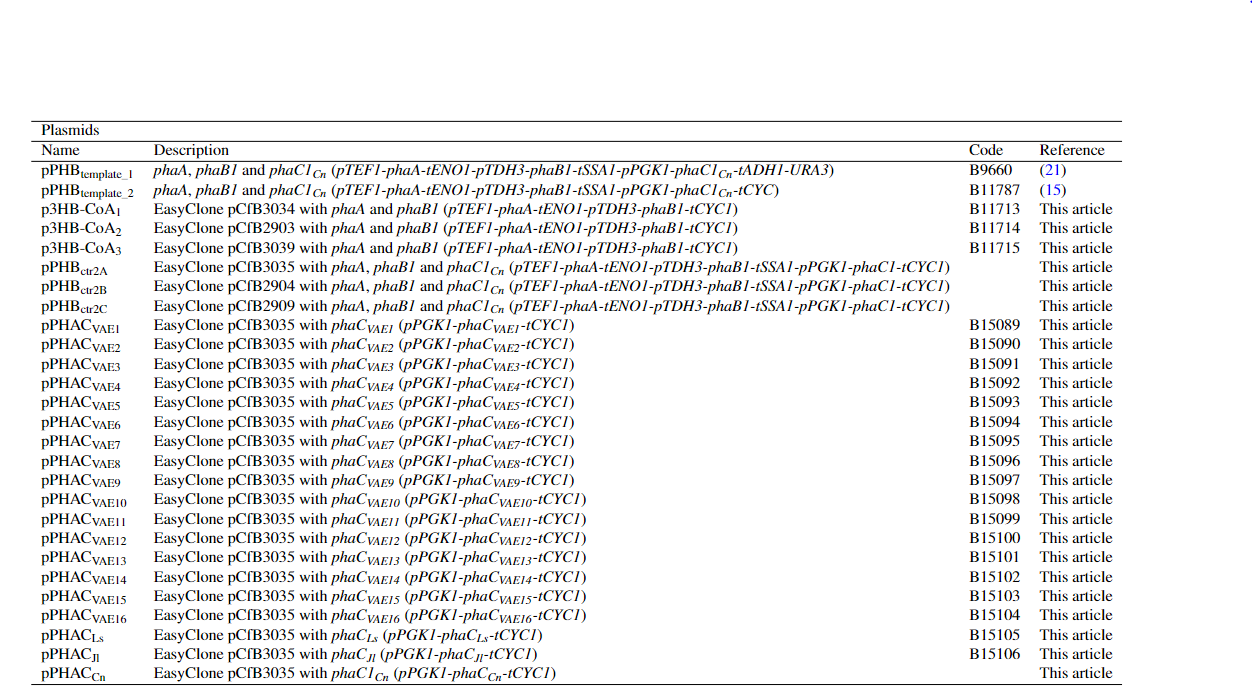
