## Supplemental figure 1 for "New-to-nature PHA synthase design using deep learning"

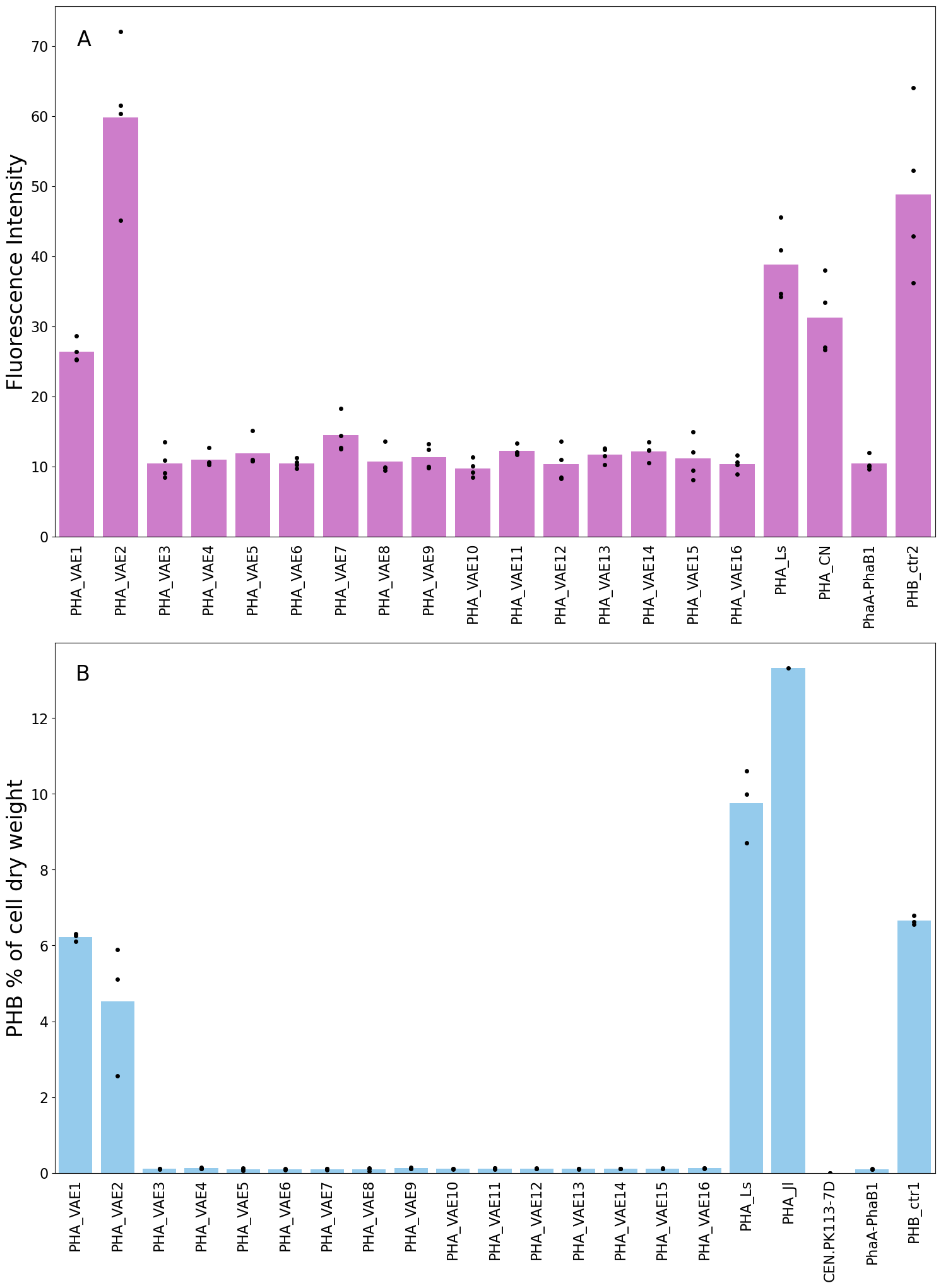

Supplementary Figure 1. Results of PHB accumulation *in vivo*. A) Fluorescence intensities of strains grown on 24-well plate and stained with Nile red method. B) The amount of PHB as % of cell dry weight quantified with GC-MS from strains grown in shake flasks. Three different biological replicates were used for strains PHA_VAE1-PHA_VAE16 and PHA_Ls. For control strains CEN.PK113-7D, PhaA-PhaB1, and PHB_ctr1 same biological replicate was cultured in three different shake flasks. For strain PHA_Jl, one biological replicate was cultivated. Bars show the average value of replicate cultures and circles values of each replicate cultures.
