## Supplemental figure 2 for "New-to-nature PHA synthase design using deep learning"

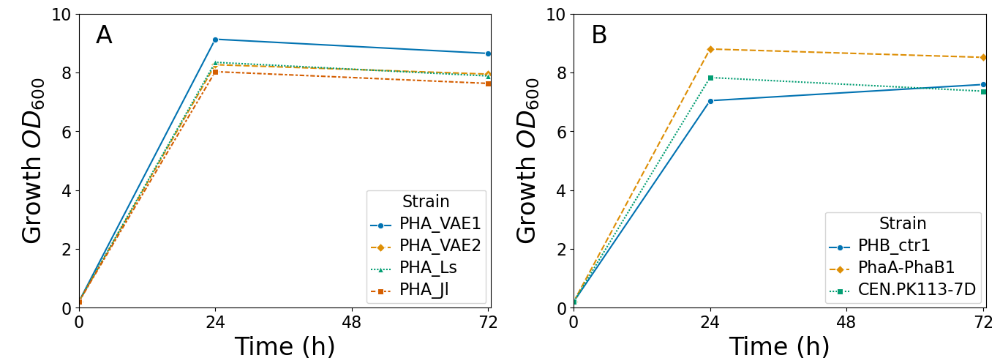

Supplementary Figure 2. Growth of the strains during the shake flask cultivation. Three biological replicates were cultivated of strains PHA_VAE1, PHA_VAE2 and PHA_Ls and one biological replicate of strain PHA_Jl. For strains PHB_ctr1, PhaA-PhaB and CEN.PK113-7D the same biological replicate was cultivated in three separate shake flasks.
