## Supplemental figure 3 for "New-to-nature PHA synthase design using deep learning"

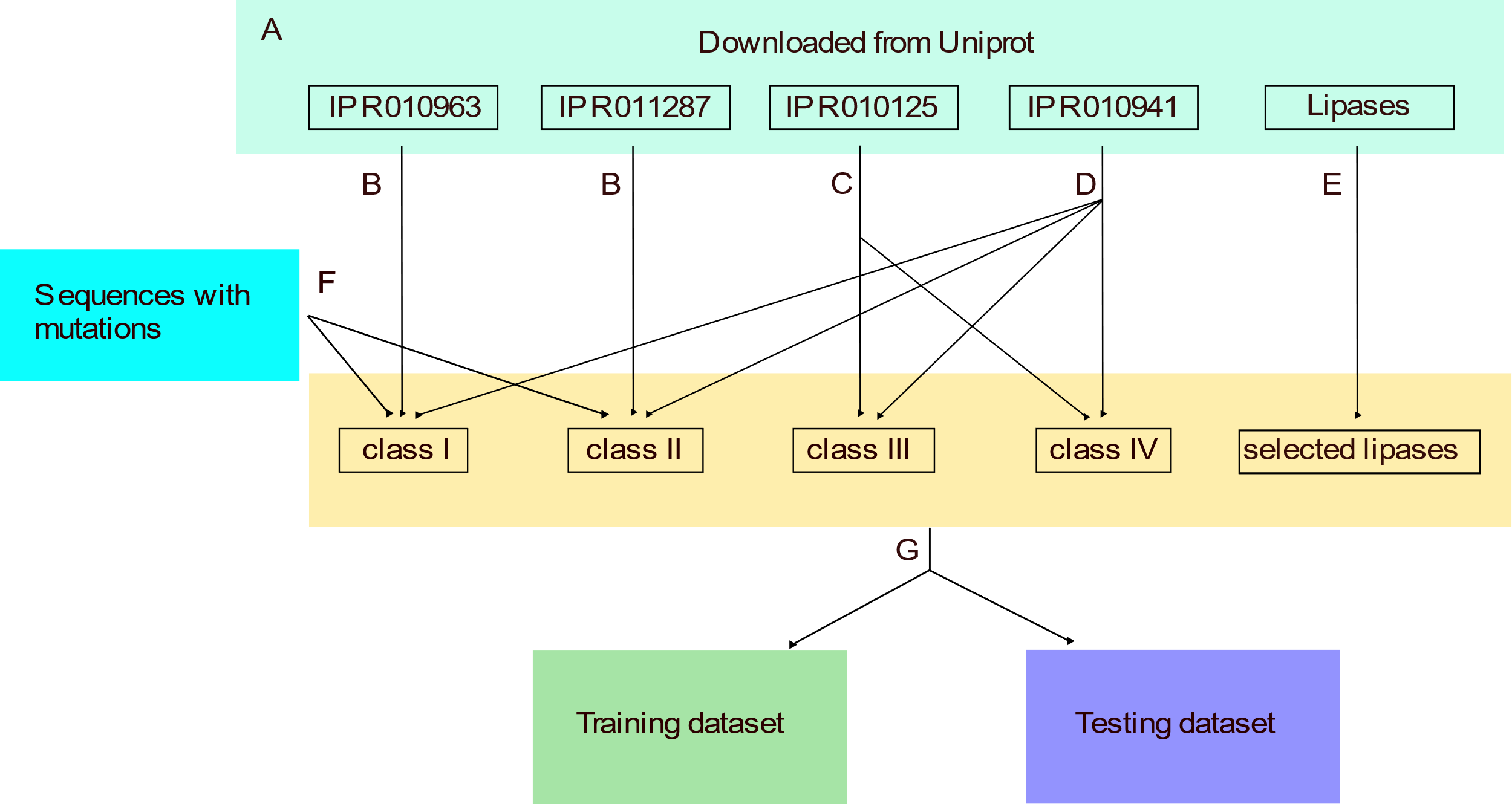

Supplementary Figure 3. Processing of training data. A) Sequences containing InterPro families IPR010963, IPR011287, IPR010125 or IPR010941 i.e. sequences containing PHA synthase class I, class II, class III or N-terminal domains were downloaded from Uniprot. In addition, sequences with lipase in their name were downloaded from Uniprot. B) Sequences of IPR010963 and IPR011287 were assigned to class I and class II respectively. C) Sequences of IPR010125 were assigned to class III or class IV based on multiple sequence alignment (MSA) and phylogenetic tree. MSA was made using ClustalW 2.1 and D3 javascript library was used to generate a phylogenetic tree from the MSA. Known class IV PHA synthase sequences (Q8GI81 and Q9ZF92, Uniprot) belonged to one cluster of sequences in the phylogenetic tree. Sequences in this cluster were assigned to class IV and the rest of the sequences in IPR010125 to class III. D) Sequences belonging to IPR010941 were assigned to classes I-IV. BLAST was run for the sequences against sequences belonging to IPR010963, IPR011287 or IPR010125. The query sequence was assigned to the same class as the hit with lowest e-value. E) BLAST was run for all lipase sequences downloaded from Uniprot against PHA synthase sequences in IPR010963, IPR011287 or IPR010125. 5000 lipase sequences with lowest e-value were selected to the dataset. F) 32 sequences with known good mutations (Supplementary Table 1) were added to classes I and II. G) The dataset was divided to training and testing data based on clustering sequences with CD-hit. Sequences of each class (class I-IV of PHA synthases and lipases) were clustered and one cluster of each class was selected as testing data. The rest of the sequences were used as training data.
